# To assemble or not to resemble – A validated Comparative Metatranscriptomics Workflow (CoMW)

**DOI:** 10.1101/642348

**Authors:** Muhammad Zohaib Anwar, Anders Lanzen, Toke Bang-Andreasen, Carsten Suhr Jacobsen

## Abstract

**Background:** Metatranscriptomics has been used widely for investigation and quantification of microbial communities’ activity in response to external stimuli. By assessing the genes expressed, metatranscriptomics provide an understanding of the interactions between different major functional guilds and the environment. Here, we present *de-novo* assembly-based Comparative Metatranscriptomics Workflow (CoMW) implemented in a modular, reproducible structure, significantly improving the annotation and quantification of metatranscriptomes. Metatranscriptomics typically utilize short sequence reads, which can either be directly aligned to external reference databases (“assembly-free approach”) or first assembled into contigs before alignment (“assembly-based approach”). We also compare CoMW (assembly-based implementation) with assembly-free alternative workflow, using simulated and real-world metatranscriptomes from Arctic and Temperate terrestrial environments. We evaluate their accuracy in precision and recall using generic and specialized hierarchical protein databases.

**Results:** CoMW provided significantly fewer false positives resulting in more precise identification and quantification of functional genes in metatranscriptomes. Using the comprehensive database M5nr, the assembly-based approach identified genes with only 0.6% false positives at thresholds ranging from inclusive to stringent compared to the assembly-free approach yielding up to 15% false positives. Using specialized databases (Carbohydrate Active-enzyme and Nitrogen Cycle), the assembly-based approach identified and quantified genes with 3-5x less false positives. We also evaluated the impact of both approaches on real-world datasets.

**Conclusions:** We present an open source *de-novo* assembly-based Comparative Metatranscriptomics Workflow (CoMW). Our benchmarking findings support the argument of assembling short reads into contigs before alignment to a reference database, since this provides higher precision and minimizes false positives.

## 1 Introduction

Metatranscriptomics provides an unprecedented insight to complex functional dynamics of microbial communities in various environments. The method has been applied to study the microbial activity in thawing permafrost and the related biogeochemical mechanisms contributing to greenhouse gas emissions [1], and Gonzalez *et al.* [2] applied metatranscriptomics to evaluate root microbiome response to soil contamination. Metatranscriptomics has also been used to study the functional human gut microbiota [3,4]. The method is typically used to identify, quantify and compare the functional response of microbial communities in natural habitats or in relation to environmental or physio-chemical impacts.

Using high-throughput sequencing techniques such as Illumina, metatranscriptomics offers a non PCR biased method for looking at transcriptional activity occurring within a complex and diverse microbial population at a specific point in time [5]. However, curation and annotation of this complex data has emerged as a major challenge. To date, several studies have used various analytic workflows. Typically, short sequence reads are utilized, which can either be individually aligned directly to external reference databases (hereafter “assembly-free”) or assembled into longer contiguous fragments (contigs) for alignment (hereafter “assembly-based”). Various studies have used either of these two general approaches. For example, Poulsen *et al.* [6] used an assembly-based approach. An open-source pipeline, IMP [7] also uses this approach in integrated metagenomic and metatranscriptomic analyses. The assembly-free Approach has instead been used by e.g. Jung *et al.* [8], aligning short reads to reference genomes of lactic acid bacterial strains associated with the kimchi microbial community. Similarly, an open source pipeline developed by Martinez *et al.* [9] to analyse metatranscriptomics data-sets also aligns short reads directly to a protein database before annotation. The choice of either of these two alternatives for metatranscriptomics analyses may depend on lack of thorough comparisons. Since no independent and direct comparison between them has been performed presently, various metatranscriptomics analysis approaches may at times produce inconsistent observations, even if identical databases are used in the analysis. Thus, standardization of computational analysis is necessary to enable further propagation of metatranscriptomics approaches and their integration into microbial ecology research. Benchmarking provides a critical view of the efficiency and precision of different workflows and use of simulated communities for benchmarking enables the analysis to be independent of experimental variation and biases [10].

Here, we present Comparative Metatranscriptomic Workflow (CoMW) implemented using the *de-novo* assembly-based approach, standardized and validated for functional annotation and quantitative expression analysis. We validated the suitability of CoMW for functional analysis by comparing it to a typical assembly-free approach using simulated datasets and evaluated the accuracy of both approaches using precision, recall and False Discovery Rates (FDR). Three different protein databases were selected for this benchmarking in order to include a representative selection of three different degrees of specialization, on a range from a more inclusive database with wide coverage (universality) and low degree of expert curation, to a smaller, highly curated database, with more narrow coverage: 1) M5nr [11] an inclusive and comprehensive non-redundant protein database in combination with eggNOG hierarchical annotation 2) Carbohydrate-Active Enzymes (CAZymes) [12] a database dedicated to describing the families of structurally-related catalytic and carbohydrate-binding modules of enzymes and 3) Nitrogen Cycling Database (NCycDB) [13] a specialized and manually curated database covering only N cycle genes. Finally, in order to estimate the consistency and variance in the results caused by the choice of approach we then applied them to real world metatranscriptomes from microbial communities in 1) active-layer permafrost soil from Svalbard [14] and 2) Ash impacted Danish Forest soil [15].

## 2 Findings

### 2.1 Comparative Metatranscriptomics Workflow (CoMW)

We have standardized, implemented, and validated a metatranscriptomic workflow (CoMW) using de-novo assembly-based approach that can assist in analysing large metatranscriptomics data. It makes each step of the metatranscriptomic workflow straightforward and help to make these complex analyses more reproducible and the components re-useable in different contexts. The core processes such as ORF detection and alignment against the functional database are vital in any metatranscriptomic analyses and are, therefore, present uniformly in all workflows. However, since most of the tools performing these core processes are ever improving, the workflow is implemented in modular format in order to have the possibility of using alternative tools and databases if preferred or use a newer version of these tools. Modularity additionally also provides choice where optional steps can be skipped, changed or even improved in a structural manner for example the scripts are designed to cater contigs from more than one assembler. In addition to core process CoMW has a couple of optional steps such as abundance based and non-coding RNA filtering which can be different in data sets from a different environment. CoMW is open source workflow written in python available at (https://github.com/anwarMZ/CoMW) and published as a computational capsule on codeocean [16]. An Anaconda cloud environment is created with the provided configuration file to install third-party tools and dependencies. Help regarding input, output and parameters is provided with each script and a comprehensive tutorial is presented in the GitHub repository.

### 2.2 Evaluation of CoMW (assembly-based Approach) and comparison to an assembly-free method

In order to compare the performance of the assembly-based workflow CoMW and assembly-free approaches, we simulated community transcript data using 4943 full length genes provided by Martinez *et al.* [9]. We analysed both approaches separately and compared against direct annotation of full-length genes. The full-length genes were annotated using all three databases (M5nr, CAZy and NCycDB) independently to classify them into functional subsystems and gene families. Figure 1 shows detailed workflow of comparative analysis using both approaches.

**Figure 1:**
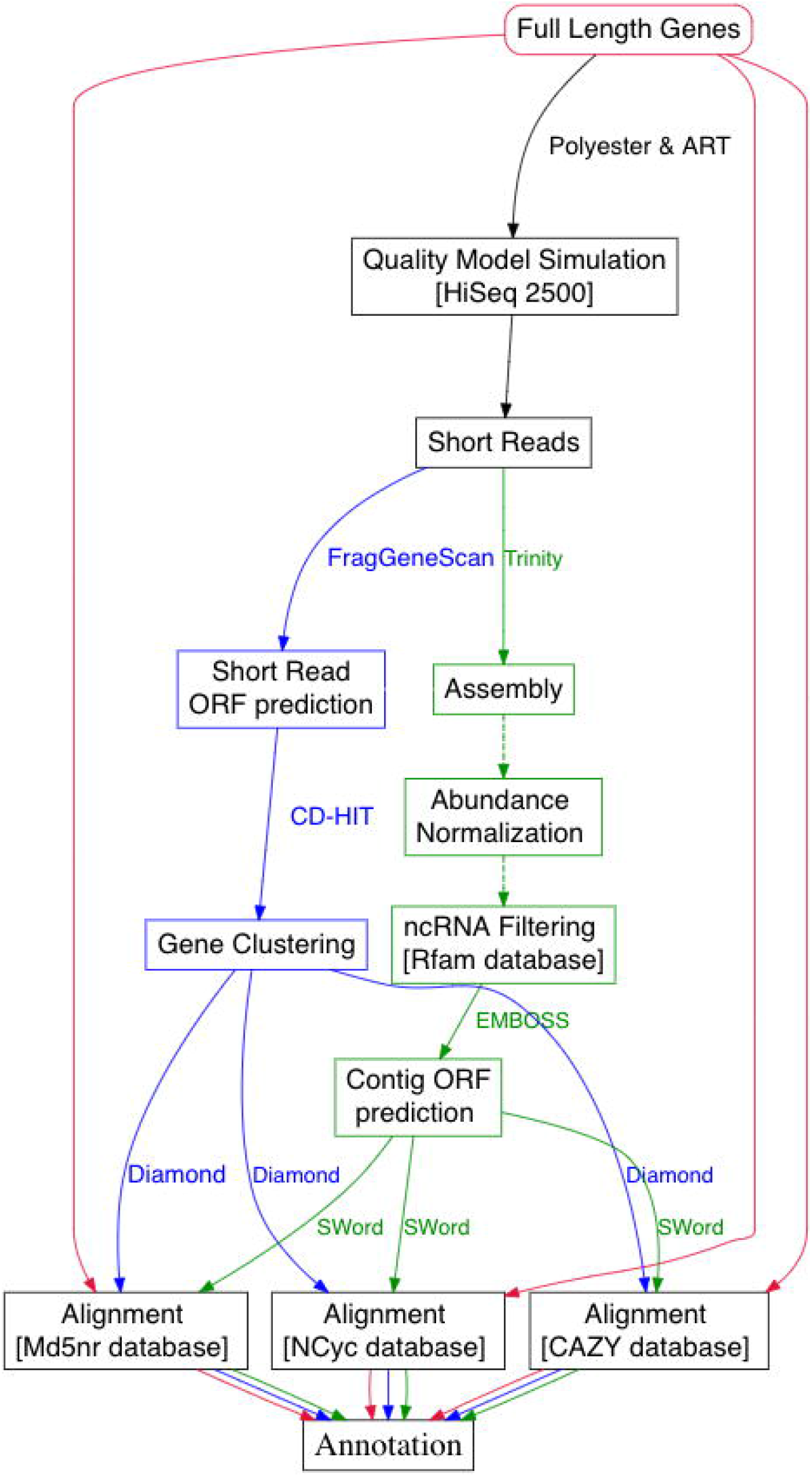
Flowchart illustrating the evaluation and benchmarking scheme used for the comparison of alternative approaches. Red path indicates the full-length genes workflow, Green indicates the steps in the assembly-based workflow CoMW and Blue indicates the steps in the assembly-free approach.

#### 2.2.1 Functional assignment

##### M5nr Alignment

Full length genes of the simulated community dataset were aligned and identified into 671 unique eggNOG orthologs, belonging to 19 distinct functional subsystems (level II). At the default confidence threshold (bit score 50), the, assembly-free approach produced alignments to 820 orthologs with a precision of 85% (14.9% FPs), whereas CoMW identified 665 orthologs with a precision of 99.3% (0.6% FPs) at the default confidence threshold of 1E-5. Repeating the alignments using a gradient of 15 varying confidence thresholds for each approach (Low – T_L_, Medium – T_M_ and High – T_H;_ five thresholds / category) resulted in dissimilar performance for both approaches. The precision and recall of CoMW did not decrease below 99.3% and 98.5% respectively throughout all categories whereas the assembly-free approach had a maximum precision of 96.3% at T_M_ and decreases to 85% at T_L_ and T_H_. CoMW also produced fewer (only 0.6%) FPs consistently compared to the assembly-free Approach of FPs ranging from 14.9% to minimum 3.6% at highest precision. Based on F-Score the most optimal alignment for each approach is given in Table 1, whereas detailed values for precision, recall, F-Score and FDR are listed in Supplementary Table S1. We then also evaluated both approaches by selectively removing sequences belonging to a certain functional subsystem from the M5nr database in a controlled manner (segmented cross validation) in order to replicate real world metatranscriptomes where a certain functional subsystem can be completely or partially absent from the reference database. We removed four (level II) subsystems (“[D] Cell cycle control, cell division, chromosome partitioning”; “[L] Replication, recombination and repair”; “[E] Amino acid transport and metabolism” and “[R] General function prediction only” and “[S] Function unknown”). The level II subsystems were randomly removed (see data availability for the script used for the removal) one at a time realigning full-length genes and simulated reads using both CoMW and assembly-free approaches to the cropped database to compare identification consistency. In each validation round, both precision and recall of CoMW were significantly higher than assembly-free approach. Recalling ability of assembly-free approach dropped significantly in this validation as compared to full database comparison. CoMW also produced less FPs as compared to assembly-free approach. Table 2 provides details for each validation cycle.

**Table 1.**
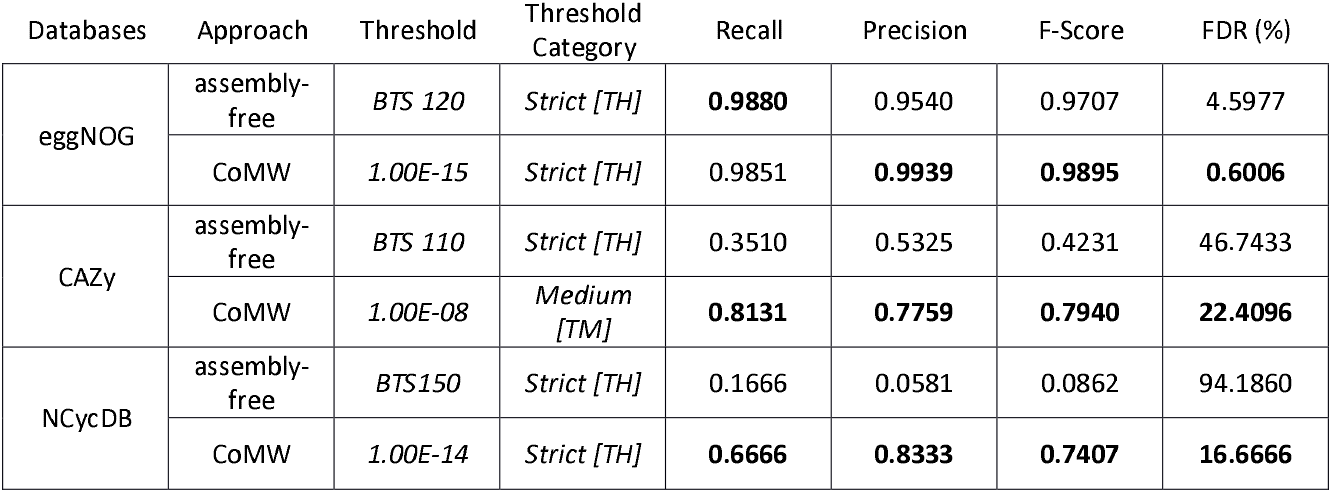
Comparison of Precision, Recall, F Score and FDR for the assembly-free and the CoMW (assembly-based) approaches using all three databases based on best F-Score (Full table for both approaches and databases can be seen in Table S1, S2 and S3). Bold emphasizes better precision, recall, F-Score and FDR in each database between both approaches

**Table 2.** Comparison of Precision, Recall, F Score and FDR for the assembly-free and CoMW (assembly-based) approaches using the selective removal of functional subsystems from eggNOG database (segmented cross-validation) to evaluate the consistency of both approaches. Bold emphasizes better consistency compared to Full length genes

| Removed Subsystem | Approach | Recall | Precision | F-Score | FDR (%) |
| --- | --- | --- | --- | --- | --- |
| Cell wall/membrane/envelope biogenesis [M] | assembly-free | 0.8726 | 0.9580 | 0.9133 | 4.1958 |
|  | CoMW | **0.9792** | **0.9855** | **0.9824** | **1.4423** |
| Replication, recombination and repair [L] | assembly-free | 0.8734 | 0.9588 | 0.9141 | 4.1166 |
|  | CoMW | **0.9796** | **0.9858** | **0.9827** | **1.415** |
| Amino acid transport and metabolism [E] | assembly-free | 0.8750 | 0.9589 | 0.9150 | 4.1095 |
|  | CoMW | **0.9812** | **0.9874** | **0.9843** | **1.2578** |
| General function prediction only and<br>Function unknown [R], [S] | assembly-free | 0.8933 | 0.9281 | 0.9104 | 7.1856 |
|  | CoMW | **0.9884** | **0.97443** | **0.9814** | **2.5568** |

##### CAZY Alignment

From 2395 full length genes, 500 sequences were aligned to 395 unique functional genes in the CAZY database, which belonged to 130 gene families and were further classified as seven enzyme classes. Using default confidence thresholds (BTS 50, 1E-5), the assembly-free approach identified 765 functional genes belonging to 112 unique families and six enzyme classes with a precision of 28.5% (71.4% FPs). CoMW identified 488 functional genes from CAZY database that were classified into 147 gene families from seven enzyme classes with a precision of 66% (FDR 33.9%) at the default confidence threshold. However, when we repeated the process with 15 various confidence thresholds, precision improved consistently and FPs decreased, whereas for the assembly-free approach, precision dropped significantly with increasing confidence threshold (see Table 1 and Supplementary Table S2).

##### NCycDB Alignment

410 out of 2395 full-length genes were aligned to this database, identified as 29 unique Nitrogen cycle genes and further belonging to 15 functional gene families in five pathways. Using default confidence thresholds, the assembly-free approach identified 1541 functional genes belonging to 25 functional gene families classified into six pathways with a precision of 0.9% (99% FPs). CoMW identified 42 Nitrogen cycle genes classified into 25 gene families from six pathways with a precision of 59.5% (40.4% FPs) at a default confidence threshold of 1E-5. Like comparisons against M5nr and CAZY we repeated the process with 15 different confidence thresholds for each approach. Precision improved significantly for CoMW at stringent thresholds whereas for the assembly-free approach, the best precision achieved was 5.8%. (Table 1, Supplementary Table S3).

#### 2.2.2 Expression Quantification

We also compared the ability of both approaches to quantify the expression of identified transcripts by performing differential expression analysis of two groups in simulated communities and compared against the full-length gene expression simulated. We selected three best identification thresholds for both approaches based on highest F-Score and performed differential expression analysis. This analysis for both approaches was carried out against all three databases using the most specific level of hierarchy in the respective databases in order to capture their ability to quantify expression levels of specific genes.

According to full-length gene alignments against eggNOG, 123 genes were significantly upregulated and 270 were significantly downregulated. According to the assembly-free Approach (with the best resulting F-Score), 73 genes were up-regulated (precision 94.5%, 5.4% FPs) and 380 (precision 65.7%, 34.2% FPs) were down regulated, whereas using the assembly-based Approach CoMW, 99 genes were identified as up-regulated (precision 94.9%, 5% FPs) and 249 down-regulated (precision 97.1%, 2.8% FPs). For the CAZy database full-length genes, 81 and 189 genes were identified as significantly up- and down regulated, respectively. Using the assembly-free approach 31 up-regulated (precision 19.3%, 80.6% FPs) and 137 down-regulated genes (precision 52.5%, 47.4% FPs) where identified, whereas the CoMW identified 83 (precision 71%, 28.9% FPs) and 191 (precision 73.8%, 26.1% Fps), respectively-In the NCyc database expression analysis, three and 14 genes were seen as significantly up and down-regulated respectively using full-length genes. According to the assembly-free approach, 26 (precision 0%, 100% FPs) and 107 (precision 4.6%, 95.3% FPs) genes were up and down regulated respectively, whereas according to CoMW, three (precision 33.3%, 66.6% FPs) genes were up-regulated and 18 (precision 55.5%, 44% FPs) were down-regulated. Precision, Recall and FDR for both approaches against all three databases are available in Supplementary Table S4. Additionally, we collapsed the functional genes into functional subsystems and gene families to remove FPs produced due to identification of homologous proteins or proteins with multiple inheritance. Fold change (log2 transformed) was then calculated for each subsystem/gene family, (see Figure 2)

**Figure 2:**
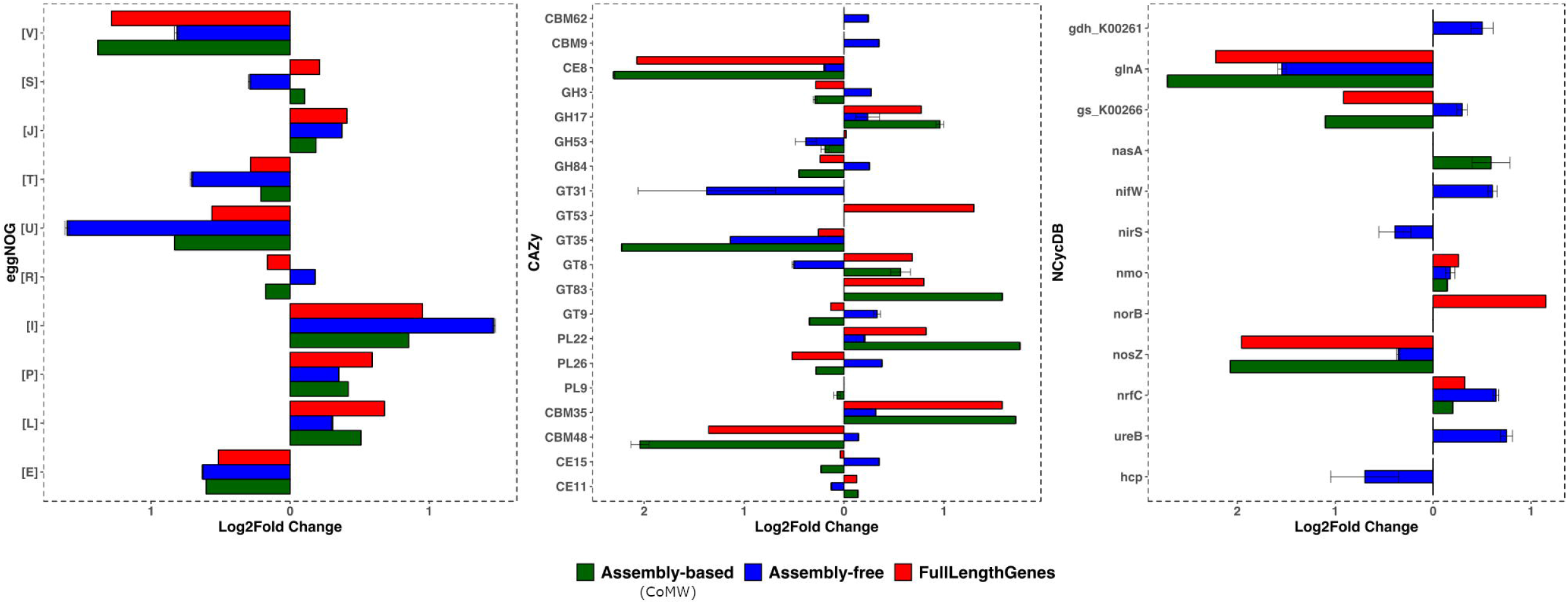
Differential Expression comparison of the assembly-free and the CoMW assembly-based approaches using A) M5nr database, B) NCycDB and C) CAZy database.

#### 2.2.3 Real-World metatranscriptomes

To evaluate the effect of the two approaches on real world data, two metatranscriptomes from microbial communities were studied. In the first study we investigated the transcriptional response during warming from −10 °C to 2 °C and subsequent cooling of 2 °C to −10 °C of an Arctic tundra active layer soil from Svalbard, Norway. The aim of the study was to understand taxonomic and functional shifts in microbial communities caused by climate change in the Arctic. A pronounced shift during the incubation period was noticed by Schostag *et al.* [14] which was not replicated by the assembly-free approach. However, using CoMW, we identified an increase of genes in the subsystem “[P] Inorganic ion transport and metabolism”. During cooling, CoMW also captured the upregulation and downregulation of genes related to “[J] Translation, ribosomal structure and biogenesis” and “[C] Energy production and conversion” respectively (Figure 3) unlike the assembly-free approach. These findings may have implications for our understanding of carbon dioxide emission, Nitrogen cycling and plant nutrient availability in Arctic soils.

**Figure 3:**
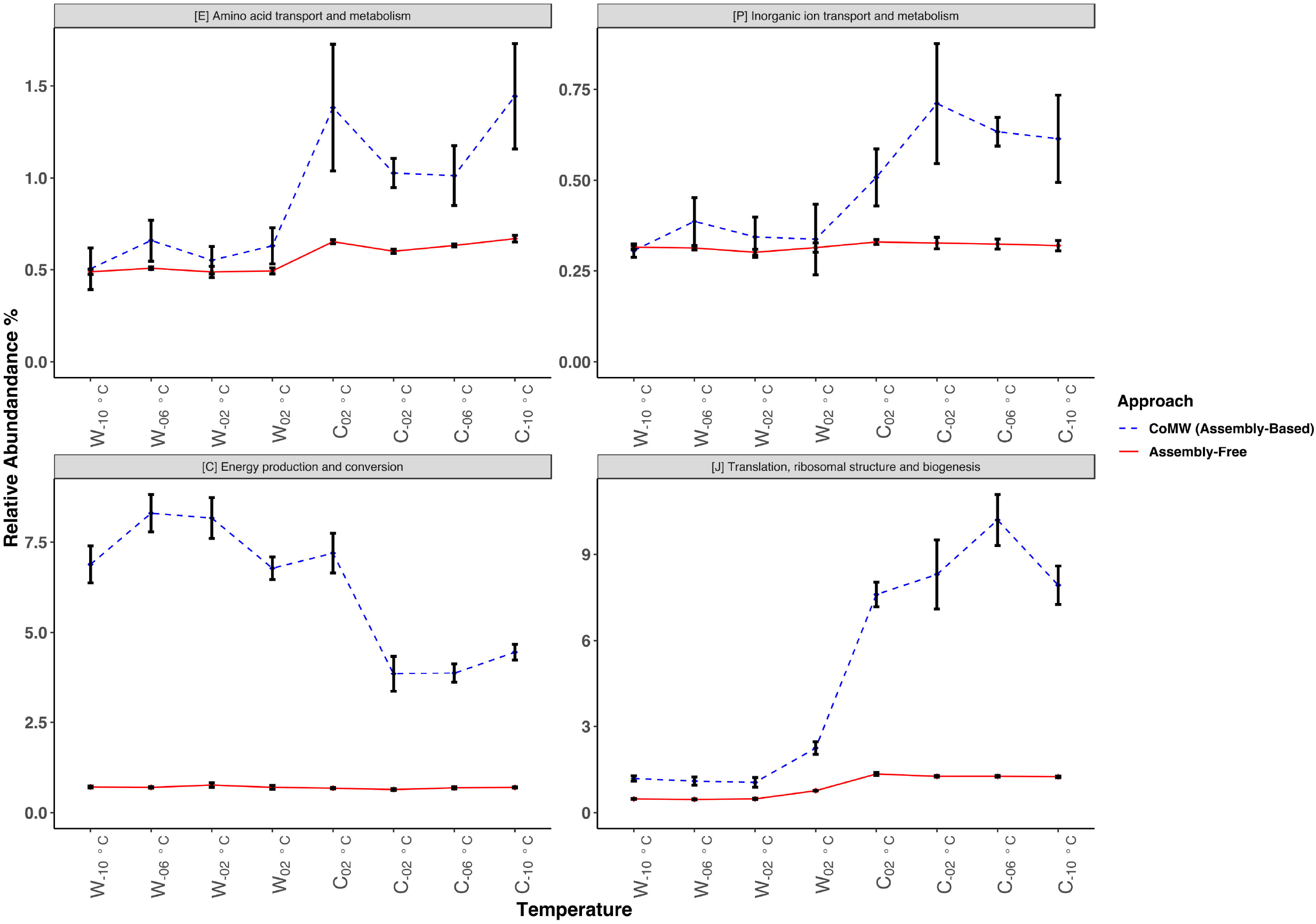
Relative abundance of eggNOG functional subsystems in Arctic permafrost soil identified and quantified using both CoMW and the assembly-free approach compares the differences in observed functional dynamics. Blue dotted line represents trends using CoMW (assembly-based) whereas Red Solid line represents assembly-free approach

In the second study, we investigated the effects of wood ash amendment on Danish forest soils [15]. Ash was added in three different quantities (0/control, 3, 12 and 90 tonnes ash per hectare (t ha^−1^)) and the effect over time was analysed in soil communities at 0, 3, 30 and 100 days after ash addition. This resulted in strong effects on functional expression as seen in Figure 4. Both approaches once again displayed varying results such as changes in genes related to eggNOG functional subsystem “[W] Extracellular structures”, assembly-free approach also identified 75% of genes as “[S] Function unknown” consistently unlike assembly-based.

**Figure 4:**
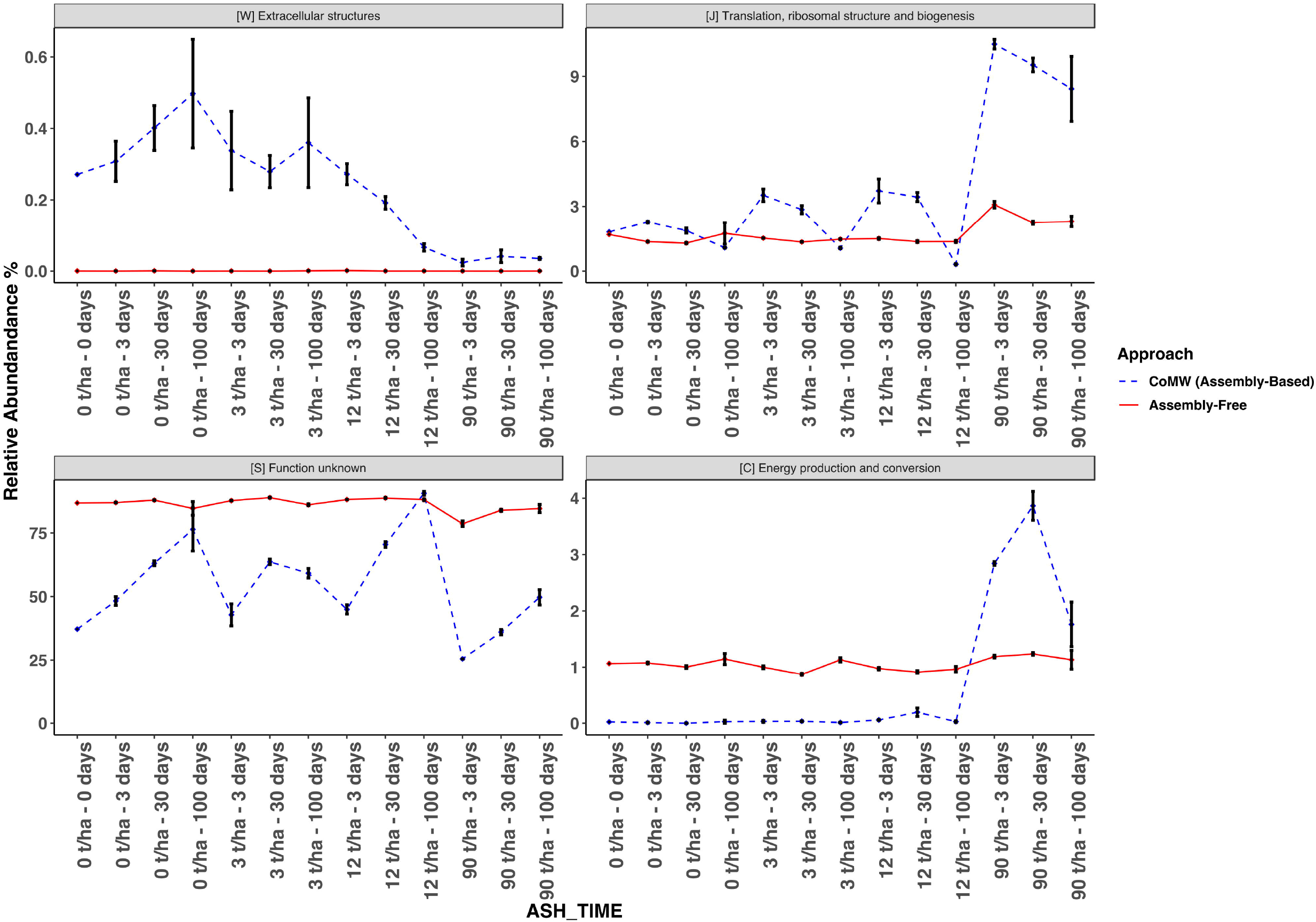
Relative abundance of eggNOG functional subsystems in Ash deposited Danish forest soil with time identified using both the CoMW and an assembly-free approach. Blue dotted line represents trends using CoMW (assembly-based) whereas Red Solid line represents assembly-free approach

## 3 Discussion

The application of metatranscriptomics is less common than other DNA-based genomics techniques and thus most analysis pipelines are built *ad hoc* [17]. An assembly-free approach is used in a few pipelines/workflows such as COMAN [18], Metatrans [9], and SAMSA2 [19], while an assembly-based approach is used in a few such as IMP [7]. The lack of thorough benchmarking studies and standardized workflows in metatranscriptomics has made it a more challenging task to analyse the typically big datasets produced. Previous studies e.g. Zhao *et al.* & Celaj *et al.* [20,21] have compared *de-novo* sequence assemblers including Trinity [22], MetaVelvet [23], Oases [24], AbySS [25] and SOAPden-ovo [26]. Similarly, for assembly-free approach direct short read mappers have been compared thoroughly such as DIAMOND [27], BLASTX [28] and RAPSearch2 [29] but an independent comparison of the two different approaches based on including assembly or directly aligning reads (here “assembly-free”) has been lacking. Critical Assessment of Metagenomic Interpreter (CAMI) [30] is so far the most comprehensive benchmarking effort, however it lacks any similar metatranscriptomics benchmarking. IMP [7] uses an integrated approach of metagenomics and metatranscriptomics and has some overlapping areas to CoMW and can be used together due to modular approach of CoMW.

Using simulated samples comprised of genes collected from abundant genomes provided by Martinez *et al.*, we show that both approaches provide similarly high recall rates against the general comprehensive database M5nr. However, CoMW provided a significantly better precision and a lower false discovery rate for identification and quantification. For relatively compact and specialized databases, recall and precision drop for both approaches (especially for the most compact database NCyc). Whereas, CoMW still appeared to be more precise, meaning that fewer genes were mis-assigned against these database and significantly lower FPs were produced.

We have attempted to assist this decision-making for processing metatranscriptomic analysis by independently assessing the performance of the two most common approaches and provide a road map for functional annotation and expression quantification against databases ranging from inclusive to specialized. The significantly higher precision in identification and quantification for gene families and functional subsystems in simulated samples, against all three databases, confirmed that while an assembly step is challenging computationally, it holds the potential to reveal information regarding the gene expressions that is not attainable without it. Selecting a single best workflow or pipeline for all types of metatranscriptomics studies is not a straightforward affair, and we believe that choice of approach changes the outcome of study significantly as observed with real-world datasets from active-layer permafrost soil from Svalbard and Ash impacted Danish Forest soil. In addition to choosing the right workflow, combining that with the appropriate reference database is equally important to ensure the best annotation performance. With databases specialized for one or more specific environments or functional categories, the assembly-free Approach under-performs due to its inability to identify alignments to homologs in the reference database. We also show that the assembly-free Approach can increase the FDR in annotation when a database is dominant in specific functional subsystem, which can also lead to wrong estimation of fold change in expression

While taxonomic annotation is beyond the scope of CoMW and thus our benchmarking analyses, it is important to consider the limited value of most functional genes for and thus functional metatranscriptomics alone for structural profiling of environmental communities, due to the high rate of horizontal gene transfer (HGT) [31]. Approaches for this purpose include the identification of a limited set of “phylogenetic marker genes” (eg.[32]) or “total RNA” metatranscriptomics whereby the rRNA content is retained and utilized for taxonomic analysis [33]. Though not shown here, we expect that the former approach would also benefit in accuracy from assembling mRNA to full length transcripts before classification, based on our results regarding functional diversity. The total RNA approach also benefits from custom rRNA targeted assembly [15], which may be incorporated into CoMW thanks to its modularity.

In summary, we present the assembly-based workflow CoMW and show that this approach results in consistently better accuracy for functional analysis of metatranscriptomics data. Our benchmarking results show that the choice of approach (assembly-free *v* assembly-based) and database significantly affects the quality of the identification, annotation and expression results. Given the impact of each of these variables, it is inevitable that it significantly affects the results of an individual study and comparison of across studies. We believe that the work presented here will both provide a useful tool for and assist the microbial ecology research community to make more informed decisions about the most appropriate methodological approach to analyze large metatranscriptomic datasets with improved precision.

## 4 Methods

### 4.1 CoMW Implementation

CoMW (assembly-based) is based on four major steps: 1) *De-novo* Assembly and Mapping; 2) Filtering; 3) Gene Prediction and Alignment 4) Annotation.

*De-novo Assembly and Mapping* of short reads back to assembled contigs is done using Trinity [22] and BWA [34] respectively. Various tools have been developed for de-novo metatranscriptome reconstruction that usually rely on graph-theory. Trinity however generates the most optimal assemblies for coding RNA reads [17,21,35]. Nevertheless, in CoMW, user can assemble short reads into contigs by any assembler preferred but it can reduce the quality of the following steps such as alignment of contigs.

*Filtering* of Contigs is done to remove variance in sequences/samples. Since CoMW is assembly- based, after we assemble the reads into longer contigs we also propose a 2-step filtering of the contigs to remove any chimeric or false contig made as a result of assembly or sequencing error by removing contigs that have an expression level less than a specific threshold and to remove any potential non-coding RNA contigs assembled. We can filter contig abundance data by removing all contigs with relative expression lower than a specific cut-off, e.g. 1% (selected based on dataset variance) of the number of sequences in the dataset with least number of sequences. This threshold is also flexible for different datasets and in some cases not required at all so CoMW allows user to bypass this step or change the threshold up and down based on data variation. The filtered contigs are subject to potential non-coding RNA filtration by aligning them against the RFam database [36] using infernal [37] which is a secondary-structure-aware aligner that predicts the secondary structure of RNA sequences and similarities based on the consensus structure models. Once again, the ncRNA filtering is an optional step in CoMW, though highly recommended in order to reduce FPs.

*Gene Prediction and Alignment* is done using Transeq from EMBOSS [38] to predict probable open reading frames (ORFs) of the contigs (customizable, by default six per contig). We used SWORD [39] as alignment tool against reference databases. SWORD can be used in parallel based on computational resources available and the aligned results are parsed and cut-off at a specific confidence threshold of combination of e-value and alignment length (usually 1e-5, can be changed given the assembly distribution in datasets).

*Annotation* of aligned transcripts from the previous step can be done using the databases such as eggNOG which is a hierarchically structured annotation using a graph-based unsupervised clustering available algorithm to produce genome wide orthology inferences. Aligned proteins are then placed into functional subsystems based on their best hits.), CAZy which is a knowledge-based resource specialized in the Glycogenomics, and NCycDB; a Nitrogen cycle database. This results in a count table with a contig and eggNOG ortholog or CAZy gene or NCyc gene having a certain count from each sample depending upon database used. This count table can be then used for differential expression using state-of-the-art expression analysis suit such as DESeq2 [40] or its wrapper SARTools [41]. For evaluation of CoMW we used the template script provided by the SARTools for DeSeq2 analysis where we specified first group of samples as the reference samples and second group as condition with a parametric mean-variance and Benjamini & Hochberg method for P adjustment [42].

### 4.2 Assembly-free Workflow

For the assembly-free approach we used the Metatrans pipeline [9], which uses FragGeneScan [43] for ORF predictions in short reads, CD-Hit [44] for gene clustering and Diamond [27] for alignment against the M5nr, CAZy and NCyc [11–13] database. We then used the same annotation script which Is included in CoMW. For expression analysis gene counts were normalized between samples using the DESeq2 [40] algorithm. Significantly differentially expressed genes were analysed in SARTools [41] using parametric relationship and p-value 0.05 as significance threshold. The Benjamini and Hochberg correction procedure [42] was used to adjust p-value. For parameters and versions of tools used in Metatrans see supplementary GitHub repository in data availability

### 4.3 Composition of Simulated Communities

In this study we utilised a set of simulated communities from Martinez *et al.* [9] where they collected 4943 genes (coding regions) from five abundant microbial genomes: *Bacteroides vulgatus* ATCC 8482, *Ruminococcus torques* L2-14, *Faecalibacterium prausnitzii* SL3/3, *Bacteroides thetaiotaomicron* VPI-5482 and *Parabacteroides distasonis* ATCC 8503. We simulated short reads into 100 samples using Polyester [45] embedded in a script provided by Martinez *et al.* [9] at coverage of 20x which resulted in a count table and short reads with 2395 genes to add the impact of sequencing coverage that the simulator mimics. The process of regulation of abundance was done by first dividing the 100 samples into two groups (“A” and “B”) and then abundance of randomly selected 10% genes was regulated up- and down up to 4- folds, in addition to this we also knocked out (0 abundance) 5% genes completely from both simulated reads and count tables. The process of selection of samples and genes was random but tracked. To include quality and coverage bias, we used the ART simulator [46] that mimics the coverage bias and thus some genes were removed to produce an equal number of reads in FASTQ format to those produced by Polyester. ART was initially trained with Hi-Seq 2500 Illumina quality error model from dataset discussed above to have a consistent error bias. After simulating FASTQ files we then extracted the quality data and bound it to the FASTA files generating new FASTQ files. With the coverage bias and quality training included we had a total of 62,035,912 reads (310,179 ± 3,454 reads/sample).

### 4.4 Evaluation Measures

We used the standard measures of precision (also named positive predictive value, PPV), accounting for how many annotations and identifications of significantly differentially expressed gene families and subsystems are correct and defined as 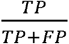 and recall (also named sensitivity or true positive rate, TPR), accounting for how many correct annotations are selected, defined as 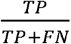 where TP indicates the number of orthologs that have been correctly annotated, FN indicates the number of orthologs/genes/functional subsystem which are in the simulated communities but were not found by a certain approach and FP indicates the number of orthologs/genes/functional subsystem that have been wrongly annotated (because they do not appear in the simulated communities). The F-score is the harmonic mean of precision and recall, defined as 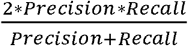.

#### Availability of source code and requirements

- Project name: Comparative Metatranscriptomics Workflow *{CoMW*)
- Project home page: https://github.com/anwarMZ/CoMW
- Operating system(s): Platform independent
- Programming language: Python, R, and bash
- Other requirements: Requirements mentioned in detailed manual at GitHub
- License: GNU General Public License v3.0

#### Availability of supporting data and materials

- Raw sequence data generated using simulation of full-length genes were deposited in the NCBI Sequence Read Archive and are accessible through BioProject accession number PRJNA509064
- Project supplementary scripts: https://github.com/anwarMZ/CoMW_supp
- Supplementary File 1 – Precision Recall Analysis of both approaches
- Supplementary File 2 – Differential Expression Analysis of all approaches using eggNOG database
- Supplementary File 3 – Differential Expression Analysis of all approaches using CAZy database
- Supplementary File 4 – Differential Expression Analysis of all approaches using NCyc database

#### Tracking and Reproducibility

- CoMW is published as computational capsule on codeocean and can be accessed through https://doi.org/10.24433/CO.1793842.v1
- CoMW is registered at SciCrunch.org with RRID – SCR_017109.

## Supporting information

Supplementary Datasheet 1

Supplementary Datasheet 2

Supplementary Datasheet 3

Supplementary Datasheet 4

## List of abbreviations

FDR: False Discovery Rate,
FP: False Positives,
TP: True Positives,
FN: False Negatives,
mRNA: messenger RNA

## Ethical Approval

Not applicable

## Consent for publication

Not applicable

## Author’s Contributions

MZA & CSJ conceived and designed the study. MZA, TBA and AL carried out the data production. MZA and AL carried out analysis. MZA drafted the manuscript and AL, TBA and CSJ revised and approved the final version.

## Acknowledgements

Authors would like to acknowledge European Commission’s MicroArctic project for the funding. We would also like to thank authors of Metatrans for providing the data used for simulation. Additionally, we would like to thank Robert Vaser author of Sword to make it available on anaconda cloud and helping in integration with CoMW.

