## Supplementary Datasheet 1 for "To assemble or not to resemble – A validated Comparative Metatranscriptomics Workflow (CoMW)"

Supplementary Table 1 Precision, Recall, F-Score and FDR for both approaches at 15 different alignment confidence thresholds using eggNOG 4.1.

| **Approach** | **Threshold** | **Threshold Category** | **Recall** | **Precision** | **F-Score** | **FDR** |
| --- | --- | --- | --- | --- | --- | --- |
| **Assembly-free approach** | BTS 10 | TL | 0.9910714 | 0.8505747 | 0.9154639 | 14.942529 |
|  | BTS 20 | TL | 0.9910714 | 0.8505747 | 0.9154639 | 14.942529 |
|  | BTS 30 | TL | 0.9910714 | 0.8505747 | 0.9154639 | 14.942529 |
|  | BTS 40 | TL | 0.9910714 | 0.8505747 | 0.9154639 | 14.942529 |
|  | BTS 50 | TL | 0.9910714 | 0.8505747 | 0.9154639 | 14.942529 |
|  | BTS 60 | TM | 0.9910714 | 0.8671875 | 0.925 | 13.28125 |
|  | BTS 70 | TM | 0.9910714 | 0.8821192 | 0.9334268 | 11.788079 |
|  | BTS 80 | TM | 0.9910714 | 0.902439 | 0.9446809 | 9.756098 |
|  | BTS 90 | TM | 0.9910714 | 0.9160935 | 0.9521086 | 8.390646 |
|  | BTS 100 | TM | 0.9895833 | 0.9366197 | 0.9623734 | 6.338028 |
|  | BTS 110 | TH | 0.9880952 | 0.9458689 | 0.9665211 | 5.413105 |
|  | BTS 120 | TH | 0.9880952 | 0.954023 | 0.9707602 | 4.597701 |
|  | BTS 130 | TH | 0.983631 | 0.9565847 | 0.9699193 | 4.341534 |
|  | BTS 140 | TH | 0.9806548 | 0.9606414 | 0.9705449 | 3.93586 |
|  | BTS 150 | TH | 0.9776786 | 0.9633431 | 0.9704579 | 3.665689 |
| **CoMW (assembly-based) approach** | 1e-1 | TL | 0.985119 | 0.993994 | 0.9895366 | 0.6006006 |
|  | 1e-2 | TL | 0.985119 | 0.993994 | 0.9895366 | 0.6006006 |
|  | 1e-3 | TL | 0.985119 | 0.993994 | 0.9895366 | 0.6006006 |
|  | 1e-4 | TL | 0.985119 | 0.993994 | 0.9895366 | 0.6006006 |
|  | 1e-5 | TL | 0.985119 | 0.993994 | 0.9895366 | 0.6006006 |
|  | 1e-6 | TM | 0.985119 | 0.993994 | 0.9895366 | 0.6006006 |
|  | 1e-7 | TM | 0.985119 | 0.993994 | 0.9895366 | 0.6006006 |
|  | 1e-8 | TM | 0.985119 | 0.993994 | 0.9895366 | 0.6006006 |
|  | 1e-9 | TM | 0.985119 | 0.993994 | 0.9895366 | 0.6006006 |
|  | 1e-10 | TM | 0.985119 | 0.993994 | 0.9895366 | 0.6006006 |
|  | 1e-11 | TH | 0.985119 | 0.993994 | 0.9895366 | 0.6006006 |
|  | 1e-12 | TH | 0.985119 | 0.993994 | 0.9895366 | 0.6006006 |
|  | 1e-13 | TH | 0.985119 | 0.993994 | 0.9895366 | 0.6006006 |
|  | 1e-14 | TH | 0.985119 | 0.993994 | 0.9895366 | 0.6006006 |
|  | 1e-15 | TH | 0.985119 | 0.993994 | 0.9895366 | 0.6006006 |

Supplementary Table 2 Precision, Recall, F-Score and FDR for both approaches at 15 different alignment confidence thresholds using CAZy database

| **Approach** | **Threshold** | **Threshold Category** | **Recall** | **Precision** | **F-Score** | **FDR** |
| --- | --- | --- | --- | --- | --- | --- |
| **Assembly-free approach** | BTS 10 | TL | 0.5707071 | 0.2684086 | 0.365105 | 73.15914 |
|  | BTS 20 | TL | 0.5707071 | 0.2684086 | 0.365105 | 73.15914 |
|  | BTS 30 | TL | 0.5707071 | 0.2684086 | 0.365105 | 73.15914 |
|  | BTS 40 | TL | 0.5707071 | 0.2684086 | 0.365105 | 73.15914 |
|  | BTS 50 | TL | 0.5505051 | 0.2853403 | 0.375862 | 71.46597 |
|  | BTS 60 | TM | 0.5176768 | 0.3410982 | 0.411234 | 65.89018 |
|  | BTS 70 | TM | 0.4621212 | 0.3567251 | 0.40264 | 64.32749 |
|  | BTS 80 | TM | 0.4292929 | 0.3846154 | 0.405728 | 61.53846 |
|  | BTS 90 | TM | 0.3964646 | 0.3974684 | 0.396966 | 60.25316 |
|  | BTS 100 | TM | 0.3787879 | 0.433526 | 0.404313 | 56.6474 |
|  | BTS 110 | TH | 0.3510101 | 0.532567 | 0.423136 | 46.7433 |
|  | BTS 120 | TH | 0.3282828 | 0.5777778 | 0.41868 | 42.22222 |
|  | BTS 130 | TH | 0.3232323 | 0.5925926 | 0.418301 | 40.74074 |
|  | BTS 140 | TH | 0.3106061 | 0.6029412 | 0.41 | 39.70588 |
|  | BTS 150 | TH | 0.3030303 | 0.6122449 | 0.405405 | 38.77551 |
| **CoMW (assembly-based) approach** | 1e-1 | TL | 0.9419192 | 0.2634181 | 0.4117 | 73.65819 |
|  | 1e-2 | TL | 0.9469697 | 0.3626692 | 0.524476 | 63.73308 |
|  | 1e-3 | TL | 0.9494949 | 0.4934383 | 0.649396 | 50.65617 |
|  | 1e-4 | TL | 0.9469697 | 0.5813953 | 0.720461 | 41.86047 |
|  | 1e-5 | TL | 0.9419192 | 0.660177 | 0.776275 | 33.9823 |
|  | 1e-6 | TM | 0.8813131 | 0.6910891 | 0.774695 | 30.89109 |
|  | 1e-7 | TM | 0.8510101 | 0.7406593 | 0.792009 | 25.93407 |
|  | 1e-8 | TM | 0.8131313 | 0.7759036 | 0.794081 | 22.40964 |
|  | 1e-9 | TM | 0.7878788 | 0.7938931 | 0.790875 | 20.61069 |
|  | 1e-10 | TM | 0.7525253 | 0.8324022 | 0.790451 | 16.75978 |
|  | 1e-11 | TH | 0.7146465 | 0.8524096 | 0.777473 | 14.75904 |
|  | 1e-12 | TH | 0.7045455 | 0.8637771 | 0.776078 | 13.62229 |
|  | 1e-13 | TH | 0.6843434 | 0.8741935 | 0.767705 | 12.58065 |
|  | 1e-14 | TH | 0.6590909 | 0.8817568 | 0.754335 | 11.82432 |
|  | 1e-15 | TH | 0.6489899 | 0.8986014 | 0.753666 | 10.13986 |

Supplementary Table 3 Precision, Recall, F-Score and FDR for both approaches at 15 different alignment confidence thresholds using NCyc database

| **Approach** | **Threshold** | **Threshold Category** | **Recall** | **Precision** | **F-Score** | **FDR** |
| --- | --- | --- | --- | --- | --- | --- |
| **Assembly-free approach** | BTS 10 | TL | 0.5 | 0.008104 | 0.015949 | 99.18963 |
|  | BTS 20 | TL | 0.5 | 0.008104 | 0.015949 | 99.18963 |
|  | BTS 30 | TL | 0.5 | 0.008104 | 0.015949 | 99.18963 |
|  | BTS 40 | TL | 0.5 | 0.008104 | 0.015949 | 99.18963 |
|  | BTS 50 | TL | 0.466667 | 0.009085 | 0.017823 | 99.0915 |
|  | BTS 60 | TM | 0.4 | 0.010093 | 0.019688 | 98.99075 |
|  | BTS 70 | TM | 0.366667 | 0.011677 | 0.022634 | 98.83227 |
|  | BTS 80 | TM | 0.333333 | 0.013755 | 0.02642 | 98.62448 |
|  | BTS 90 | TM | 0.266667 | 0.014925 | 0.028269 | 98.50746 |
|  | BTS 100 | TM | 0.2 | 0.015625 | 0.028986 | 98.4375 |
|  | BTS 110 | TH | 0.2 | 0.020761 | 0.037618 | 97.92388 |
|  | BTS 120 | TH | 0.2 | 0.026316 | 0.046512 | 97.36842 |
|  | BTS 130 | TH | 0.2 | 0.032787 | 0.056338 | 96.72131 |
|  | BTS 140 | TH | 0.166667 | 0.04065 | 0.065359 | 95.93496 |
|  | BTS 150 | TH | 0.166667 | 0.05814 | 0.086207 | 94.18605 |
| **CoMW (assembly-based) approach** | 1e-1 | TL | 0.833333 | 0.09434 | 0.169492 | 90.56604 |
|  | 1e-2 | TL | 0.833333 | 0.19685 | 0.318471 | 80.31496 |
|  | 1e-3 | TL | 0.833333 | 0.373134 | 0.515464 | 62.68657 |
|  | 1e-4 | TL | 0.833333 | 0.5 | 0.625 | 50 |
|  | 1e-5 | TL | 0.833333 | 0.595238 | 0.694444 | 40.47619 |
|  | 1e-6 | TM | 0.7 | 0.6 | 0.646154 | 40 |
|  | 1e-7 | TM | 0.7 | 0.65625 | 0.677419 | 34.375 |
|  | 1e-8 | TM | 0.7 | 0.677419 | 0.688525 | 32.25806 |
|  | 1e-9 | TM | 0.7 | 0.724138 | 0.711864 | 27.58621 |
|  | 1e-10 | TM | 0.7 | 0.75 | 0.724138 | 25 |
|  | 1e-11 | TH | 0.7 | 0.807692 | 0.75 | 19.23077 |
|  | 1e-12 | TH | 0.666667 | 0.8 | 0.727273 | 20 |
|  | 1e-13 | TH | 0.666667 | 0.8 | 0.727273 | 20 |
|  | 1e-14 | TH | 0.666667 | 0.833333 | 0.740741 | 16.66667 |
|  | 1e-15 | TH | 0.633333 | 0.826087 | 0.716981 | 17.3913 |

Supplementary Table 4 Precision, Recall, F-Score and FDR of significantly expressed genes for both approaches against all 3 databases based on highest F-Score from Supplementary tables 1,2 & 3

| **DB** | **Approach** | **Threshold** | **Regulation** | **Recall** | **Precision** | **F score** | **FDR** |
| --- | --- | --- | --- | --- | --- | --- | --- |
| **M5nr** | CoMW (assembly-based) | 1e-15 | down | 0.896296 | 0.971888 | 0.932563 | 2.811245 |
|  | CoMW (assembly-based) | 1e-15 | up | 0.7642276 | 0.9494949 | 0.8468468 | 5.050505 |
|  | Assembly-free | BTS 120 | down | 0.9185185 | 0.6908078 | 0.7885533 | 30.91922 |
|  | Assembly-free | BTS 120 | up | 0.601626 | 0.9487179 | 0.7363184 | 5.128205 |
|  | Assembly-free | BTS 120 | down | 0.9259259 | 0.6578947 | 0.7692308 | 34.21053 |
|  | Assembly-free | BTS 120 | up | 0.5609756 | 0.9452055 | 0.7040816 | 5.479452 |
|  | Assembly-free | BTS 120 | down | 0.9222222 | 0.6784741 | 0.7817896 | 32.15259 |
|  | Assembly-free | BTS 120 | up | 0.5609756 | 0.9583333 | 0.7076923 | 4.166667 |
| **CAZy** | CoMW (assembly-based) | 1e-7 | down | 0.78307 | 0.76684 | 0.77487 | 23.31606 |
|  | CoMW (assembly-based) | 1e-7 | up | 0.7284 | 0.70238 | 0.71515 | 29.7619 |
|  | CoMW (assembly-based) | 1e-8 | down | 0.74603 | 0.73822 | 0.74211 | 26.17801 |
|  | CoMW (assembly-based) | 1e-8 | up | 0.7284 | 0.71084 | 0.71951 | 28.91566 |
|  | CoMW (assembly-based) | 1e-9 | down | 0.74074 | 0.72539 | 0.73298 | 27.46114 |
|  | CoMW (assembly-based) | 1e-9 | up | 0.71605 | 0.725 | 0.7205 | 27.5 |
|  | Assembly-free | BTS 110 | down | 0.40741 | 0.47239 | 0.4375 | 52.76074 |
|  | Assembly-free | BTS 110 | up | 0.12346 | 0.25 | 0.16529 | 75 |
|  | Assembly-free | BTS 120 | down | 0.40741 | 0.47239 | 0.4375 | 52.76074 |
|  | Assembly-free | BTS 120 | up | 0.12346 | 0.25 | 0.16529 | 75 |
|  | Assembly-free | BTS 120 | down | 0.38095 | 0.52555 | 0.44172 | 47.44526 |
|  | Assembly-free | BTS 120 | up | 0.07407 | 0.19355 | 0.10714 | 80.64516 |
| **NCycDB** | CoMW (assembly-based) | 1e-12 | down | 0.7857143 | 0.5 | 0.61111 | 50 |
|  | CoMW (assembly-based) | 1e-12 | up | 0.3333333 | 0.25 | 0.28571 | 75 |
|  | CoMW (assembly-based) | 1e-13 | down | 0.7142857 | 0.47619 | 0.57143 | 52.381 |
|  | CoMW (assembly-based) | 1e-13 | up | 0.3333333 | 0.25 | 0.28571 | 75 |
|  | CoMW (assembly-based) | 1e-14 | down | 0.7142857 | 0.55556 | 0.625 | 44.444 |
|  | CoMW (assembly-based) | 1e-14 | up | 0.3333333 | 0.33333 | 0.33333 | 66.667 |
|  | Assembly-free | BTS 130 | down | 0.3571429 | 0.0303 | 0.05587 | 96.97 |
|  | Assembly-free | BTS 130 | up | 0 | 0 | 0 | 100 |
|  | Assembly-free | BTS 140 | down | 0.3571429 | 0.04 | 0.07194 | 96 |
|  | Assembly-free | BTS 140 | up | 0 | 0 | 0 | 100 |
|  | Assembly-free | BTS 150 | down | 0.3571429 | 0.04673 | 0.08264 | 95.327 |
|  | Assembly-free | BTS 150 | up | 0 | 0 | 0 | 100 |
